# Antiviral CD4^+^ T and myeloid cell responses to influenza vaccines are attenuated in older adults

**DOI:** 10.1101/2025.04.30.651528

**Authors:** Jared M. Oakes, Joshua D. Simmons, Cindy Hager-Nochowicz, Leslie Kirk, Joan Eason, Natasha B. Halasa, H. Keipp Talbot, Samuel S. Bailin, Jessica L. Castilho, Spyros A. Kalams

## Abstract

Recent influenza vaccine formulations have improved the magnitude of B-cell antibody responses in older adults; however, older adults remain significantly at risk for severe influenza-related illness. Although antibodies are an important metric of vaccine effectiveness, they only represent one aspect of the immune response. In this study, we combined in vitro and ex vivo assays with human samples to investigate B, CD4^+^ T, and myeloid cell responses to influenza vaccine antigens. We found that older adults mounted equivalent antibody titers to younger adults but had fewer influenza-specific CD4^+^ T cells and reduced antiviral-associated T helper cell populations. Single-cell transcriptomics revealed that older adults had attenuated interferon transcriptional signatures in T helper and myeloid cell subsets. These data suggest that with aging, transcriptional programming alterations in myeloid cells contribute to reduced antiviral T cell responses, and formulating vaccines tailored to myeloid responses is necessary to improve outcomes in older adults.

## INTRODUCTION

Older adults over 65 years constitute roughly 60% of influenza-related deaths globally,^1^ and have reduced hemagglutinin (HA) antibody responses compared to younger adults when given the same standard-dose vaccine.^2–4^ Improved vaccine formulations, first implemented in 2010 for adults over 65, have resulted in larger fold increases and higher HA-directed antibody titers by hemagglutination inhibition (HAI).^5,6^ While three improved vaccine formulations have been approved for this age group, older adults remain disproportionately affected by influenza-related illness each year. Despite evidence that these improved vaccines marginally improve cellular responses in older adults, a thorough single-cell analysis of CD4^+^ T cell responses comparing younger and older adults has not been reported.

Vaccination is our first line of defense in mitigating infectious diseases. Unlike most vaccine-preventable communicable diseases, it is recommended in the United States that influenza vaccines be administered yearly for all individuals. The typical influenza vaccine includes two influenza A (H1N1 and H3N2) and two influenza B inactivated viral strains enriched for the surface attachment glycoprotein hemagglutinin (HA) and a minor amount of other viral proteins. Since 2010, the effectiveness of HA-centric inactivated vaccines in preventing influenza disease has varied between 19% and 60% in the United States.^7^ Measures of influenza vaccine immunogenicity for most populations rely on a minimum inhibition of the HA binding site by serum antibodies (HAI). Currently, the threshold for a protective HAI antibody titer is 40, and a successful immune response to the vaccine is defined as a >4-fold rise in HAI titer.^5,8^ However, the minimum HAI titer protection threshold may be higher for more at-risk populations.^9^

Changes to the immune system resulting from aging are multifaceted. Lymphocytes, in particular, undergo significant changes in phenotypic and molecular characteristics. Older adults have decreased frequencies of B cell precursors and increased frequencies of IgD^−^ CD27^−^ late memory and low PAX5-expressing B cells, which negatively correlate with influenza-specific antibody titers following vaccination.^10–12^ Late memory B cells, in particular, also show a reduced capacity to interact with T cells during antigen presentation and co-stimulation.^13^ High-dose vaccination elicits more antibody-secreting plasmablasts in older adults; however, aging still has a negative impact on B cell receptor sequence diversity. Notably, after vaccination, older adults have increased clonal expansion of plasmablasts that produce cross-reactive antibodies with minimal increases in affinity maturation to new antigens compared to younger adults.^14^ Furthermore, older adults have reduced frequencies of virus-specific cytokine-producing cellular responses after high-dose vaccination.^15^ These age-related immune response changes are thought to impair long-term protection and lower initial HAI titers in the subsequent year.^16^

Several features of immunological aging on the adaptive immune response have been well described. Thymic involution results in reduced naïve T cell output and expansion of terminally differentiated memory cells, thus inhibiting the ability to generate a *de novo* T cell-mediated response.^17,18^ With age, memory CD4^+^ T cells exhibit reduced T cell receptor diversity, and T helper responses shift away from antiviral Th1 phenotypes with increases in IL4 and IL17 cytokine secretion. This results in fewer antigens recognized by memory T cells and effector responses not adept at combating viral infections.^19–24^ Despite these deficiencies, improved vaccine formulations elicit minor improvements in cellular immune responses measured by cytokine secretion in older adults.^25^ Research on influenza vaccination, aging, and T cells has primarily focused on circulating T follicular helper (cTfh) cells because of their role in promoting the maturation of antibody responses.^26^ While antibody responses do not correlate with cTfh responses in older adults receiving the standard-dose vaccine, high-dose vaccine formulations for older adults improve cTfh activation, which significantly correlates with HAI titer one month post-vaccination.^27,28^ Transcriptional programming of cTfh cells in older adults shows increased expression of proinflammatory STAT5- and TNF-associated gene sets linked to poor antibody responses.^29,30^ Although aging and cytomegalovirus (CMV) infection have been strongly correlated, the role of CMV in a T-cell-based immune response remains debated. There is conflicting evidence regarding the effect of CMV infection on influenza vaccine responses. Two studies have shown opposite effects where CMV correlated with better or worse vaccine responses in healthy individuals measured by influenza-specific antibodies, CD8^+^ T cell responses, or circulating cytokines.^31,32^

The impact of aging on myeloid cells and vaccine responses is less well understood compared to lymphoid cells. With age, there is an increase in myelopoiesis, primarily manifesting as an increase in circulating monocytes.^33,34^ Despite this, circulating monocytes in older adults show few transcriptional differences compared to those in younger adults.^22,35,36^ However, dendritic cells from older adults show a decline in effector functions necessary for interaction with other immune cells and cell activation through pattern recognition receptors.^37^

In this study, we examined the effect of aging on broader CD4^+^ T cell populations following influenza vaccination in adults under 45 and over 65 years of age. We coupled primary human samples with high-dimensional cytometric approaches and large-scale single-cell transcriptomics to examine molecular and phenotypic changes in CD4^+^ T cells. Older adults from this cohort exhibited robust antibody responses; however, CD4^+^ T cell responses were significantly lower compared to those of younger adults. The transcriptional programming of CD4+ T and myeloid cells in older adults revealed diminished interferon responses compared to younger adults. These findings highlight new potential targets for improved vaccine formulations in older adults.

## RESULTS

### Older and younger adults mount robust antibody responses to influenza vaccine antigens

Details of our cohort and experimental workflow are shown in Fig. 1A. We first assessed HAI antibody titers before and after vaccination. Both older and younger adults responded to vaccination with significantly increased H1N1 titers at day 7 and 28 (Fig. 1B). Younger adults had higher H1N1 titers prior to vaccination (Fig. 1C), but both age groups achieved similar titers to H1N1 by day 28 (Fig. 1D) since older adults exhibited larger fold-changes in H1N1 titers (Fig. 1E). Similarly, H3N2 titers increased significantly after vaccination within each age group (Fig. 1F). In this case, both age groups had similar H3N2 titers before vaccination (Fig. 1G), achieved similar titers at day 28 (Fig. 1H), and had a median 4-fold increase in H3N2 titer at day 28 (Fig. 1I). HAI antibody responses to the B-Victoria vaccine component followed a similar trend, with both age groups exhibiting significant increases on days 7 and 28 compared to day 0 (Fig. S1A). Older adults had a non-statistically significant difference in B-Victoria titers at day 0 and day 28 (Fig. S1B-C) and higher fold-change antibody titers against B-Victoria compared to younger adults (Fig. S1D). Only younger adults had a statistically significant increase at days 7 and 28 compared to baseline against the B-Yamagata vaccine component, while older adults had no significant change in B-Yamagata-directed titers after vaccination (Fig. S1E). Younger adults had higher B-Yamagata antibody titers before vaccination and higher titers at day 28 but no increased antibody titer compared to older adults (Fig. S1F-H). Although older adults had lower baseline HAI titers to H1N1 and B-Yamagata vaccine strains compared to younger adults, post-vaccination HAI antibody responses were comparable between age groups. Despite differences in antibody response patterns, both younger and older adults mounted serologic responses to vaccination.

**Figure 1.**
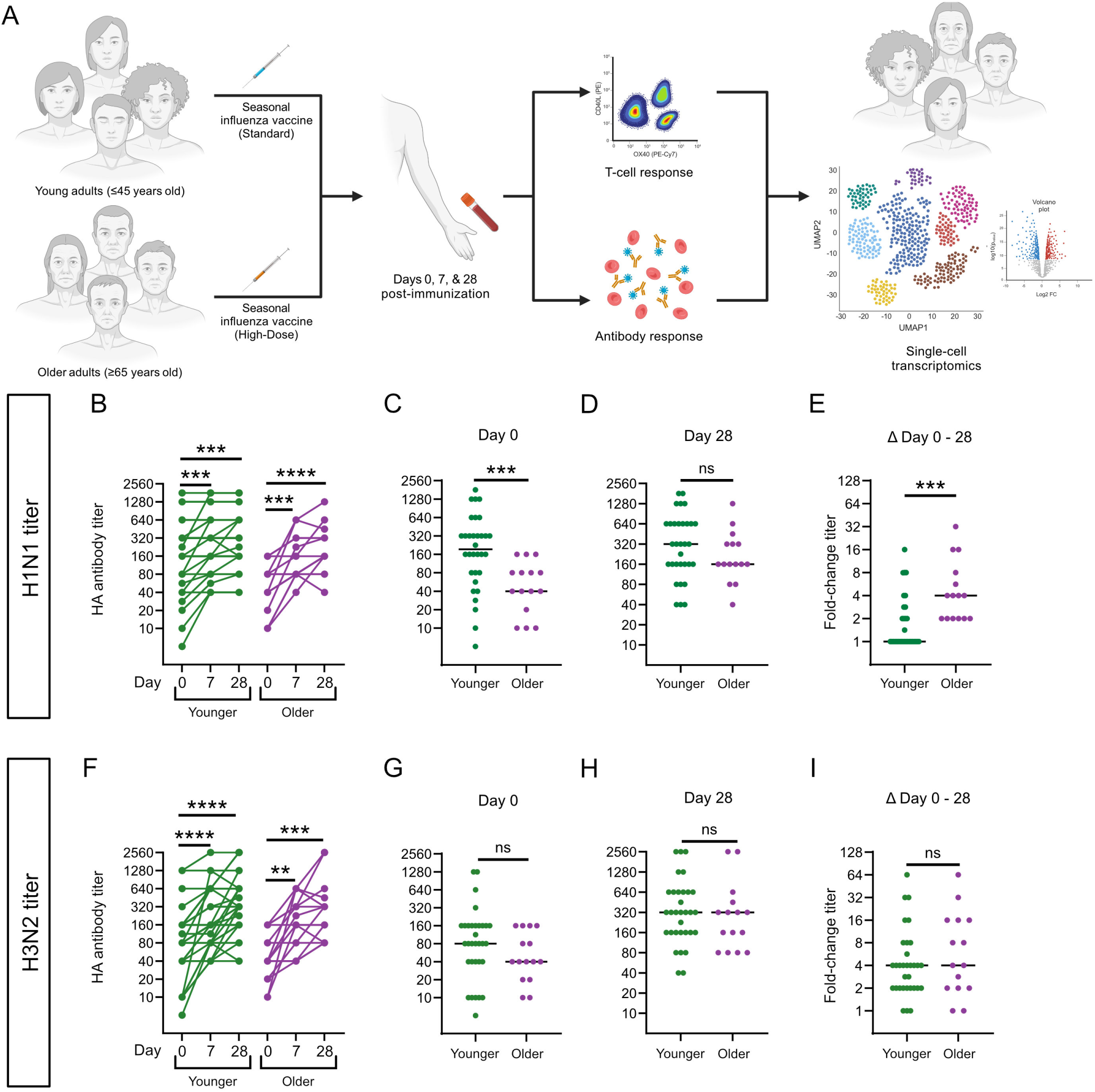
Older and younger adults mount robust antibody responses to seasonal vaccine antigens. Hemagglutination inhibition (HAI) antibody titers against H1N1 (A-D) or H3N2 (E-H) for younger (green) and older (purple) adults. (A, E) Antibody titers at days 0-, 7-, and 28-days post-vaccination with significance determined using paired Wilcoxon rank test with Bonferroni correction. (B, F) Comparison of pre-vaccination HAI antibody titers between age groups. (C, G) Endpoint HAI antibody titers at day 28 compared between younger and older adults. (D, H) Fold change in HAI antibody titer from day 0 (pre-vaccination) to day 28 (post-vaccination). (B-D, F-H) Statistics represent Mann-Whitney U test and medians of age groups are represent by a solid line. For statistical tests *p <0.05, **p <0.01, ***p <0.001, ****p <0.0001, and ns being not significant.

### Older adults lack memory CD4^+^ T cell responses to vaccine antigens

Influenza-specific CD4^+^ T cell responses were assessed using an activation-induced marker (AIM) assay. Activated cells were identified as live CD3+ CD4+ memory cells, which concurrently expressed OX40 and CD40L (Fig. S2A). Example flow cytometry plots for unstimulated (media), and influenza antigen-stimulated memory CD4^+^ T cells are shown in Fig. 2A. Upon stimulation with the 2019 H1N1 vaccine component, younger adults showed a significant increase in OX40^+^ CD40L^+^ memory CD4^+^ T cells at day 28 post-vaccination while older adults showed no change in H1N1 reactivity on day 7 or 28 post-vaccination. Only younger adults had a significant change in H3N2-reactive cells at day 7 post-vaccination (Fig. 2B). Comparing age groups, younger adults had increased frequencies of H1N1-reactive CD4^+^ memory T cells than older adults at day 0 and day 7 (Fig. 2C). There was no difference between age groups when stimulated with H3N2 prior to vaccination; however, younger adults showed more H3N2-reactive cells on day 7 post-vaccination (Fig. 2D). Despite improved vaccine formulations, older adults in our cohort have reduced influenza-specific memory CD4^+^ T cell responses compared to younger adults.

**Figure 2.**
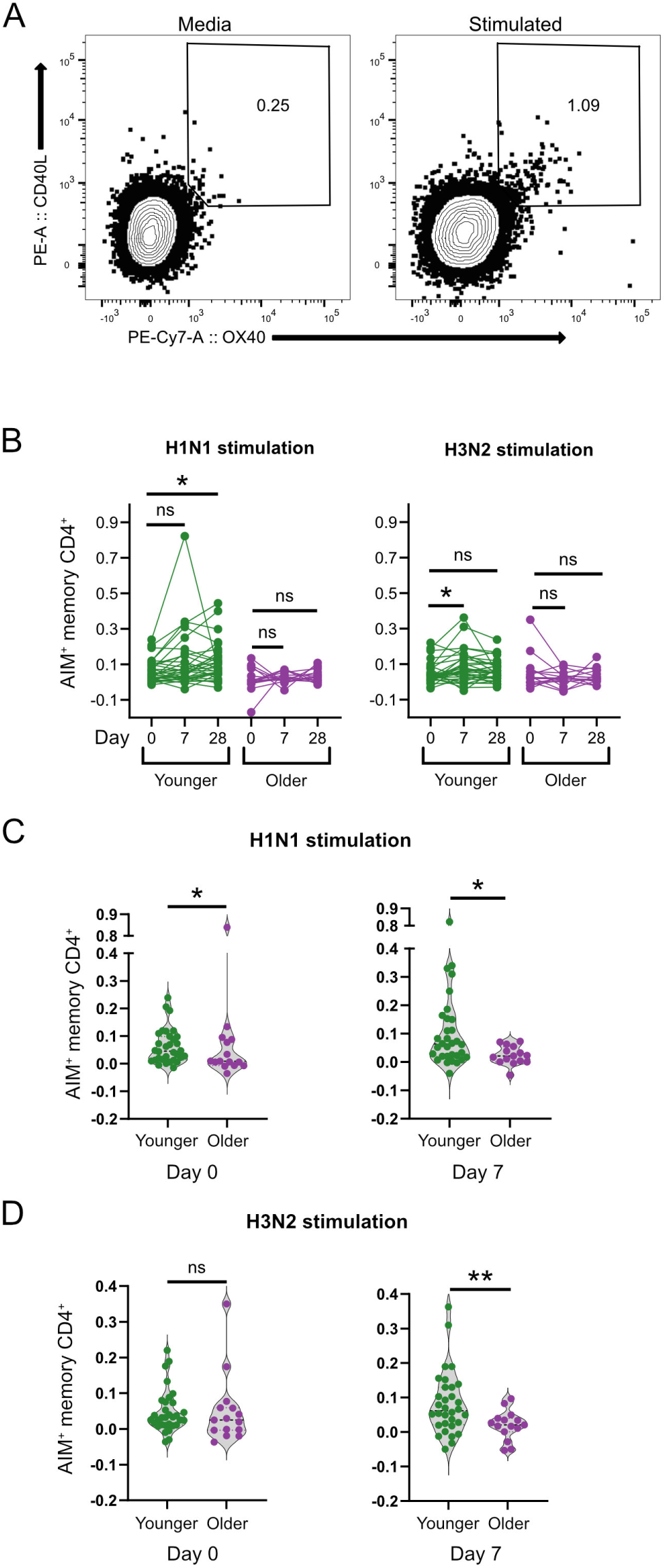
Older adults have reduced influenza-specific memory CD4^+^ T cells. (A) Example flow cytometry plots defining antigen-specific memory CD4^+^ T cells. (B) Comparison of antigen-reactive CD4+ T cells at 0-, 7-, or 28-days post-vaccination in older (purple) and younger (green) adults assessed by paired Wilcoxon rank test with Bonferroni correction. (C-D) Comparison of antigen-specific CD4^+^ T cells between older and younger adults analyzed by Mann-Whitney U test. Percentages of activated cells without stimulation have been subtracted from the stimulated conditions for each antigen. For statistical tests *p <0.05, **p <0.01, ***p <0.001, and ns being not significant.

We also investigated whether CMV infection was associated with the serologic or cellular immune responses described earlier. Fold-change in HAI antibody titers at day 28 was independent of CMV status in both age groups, with no statistically significant correlations (Fig. S3C-D). Memory CD4^+^ T cell activation assessed by AIM assay at day 7 post-vaccination also did not correlate with CMV status in either age group (Fig. S3E-F). We found no relationship between CMV seropositivity and vaccine-specific responses in either age group.

### Older adults have reduced cTfh and antiviral T helper populations not identified using unsupervised machine learning methods

Given significant differences in CD4^+^ T cell responses to the influenza vaccine antigens between younger and older adults, we used a high-parameter mass cytometry panel to phenotype PBMCs and understand compositional differences in the CD4^+^ T cell compartment. We specifically examined the differences in CD4^+^ T cell population frequencies between age groups using machine learning methods such as uniform manifold approximation and projection (UMAP) dimensionality reduction and Leiden clustering. We identified CD4^+^ T cells and used surface markers to define T cell subsets and phenotypes (Fig. 3 A-B). Overall frequencies of major cell populations, including the frequency of senescent memory CD4^+^ T cells (cluster 8), did not differ between older and younger adults (Fig. 3C). With the generated UMAP, we also compared pre-vaccination CD4^+^ T cells between age groups using Tracking Responders EXpanding (T-REX).^38^ This technique employs a k-nearest-neighbor (kNN) search, followed by a similarity threshold set at 90% for our data to identify regions significantly different between comparator groups. T-REX identified significant differences between older and younger adults in four areas on the UMAP (Fig. 3D). These populations represent effector memory (younger_1), CD57^+^ senescent (younger_2), and cTfh cells in younger adults (younger_3) and effector memory cells in older adults (older_1). However, these populations represent less than 2% of CD4^+^ T cells per individual, and are driven by select few individuals, likely due to subtle differences in surface marker expression (Fig. 3E). Since we observed no significant differences in CD4^+^ T cell population frequencies between age groups using machine learning methods, we manually identified major B cell and memory CD4^+^ T helper cell populations known to be involved in an antiviral immune response through a biaxial gating strategy (Fig. S2B). T helper subsets have been previously shown to shift from an antiviral Th1 phenotype to a Th2 and Th17 phenotype, potentially leading to improper immune activation of non-antiviral effector cells.^22–24^ We found that older and younger adults had similar frequencies of Th17 (CXCR3^−^ CCR4^+^ CCR6^+^) cells; however, older adults had decreased frequencies of antiviral Th1 (CXCR3^+^ CCR4^−^) cells and increased frequencies of extracellular antigen-targeting Th2 (CXCR3^−^ CCR4^+^ CCR6^−^) cells compared to younger adults (Fig. 3F).

**Figure 3.**
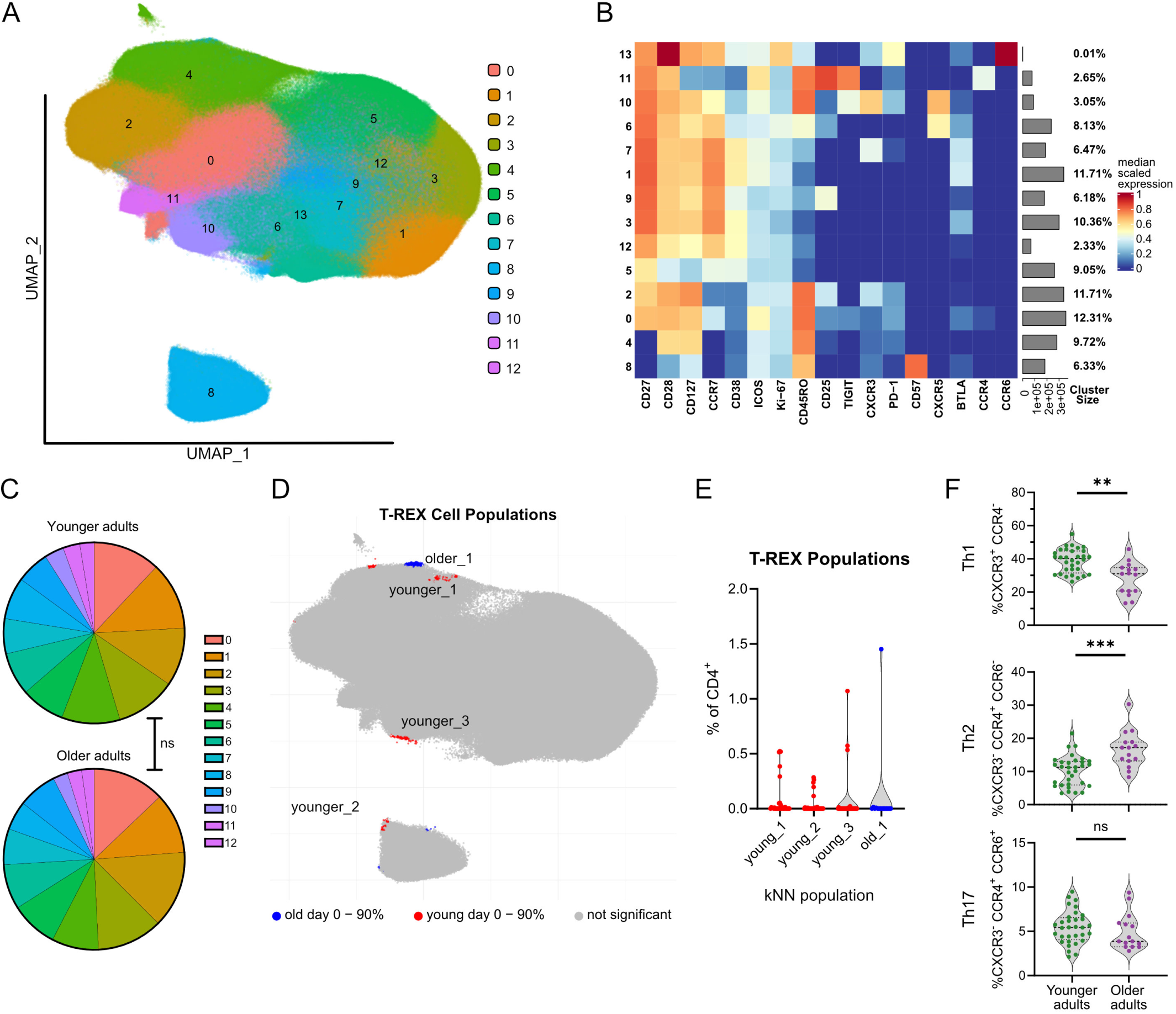
Older adults have a reduced antiviral T helper cell phenotype not identified using machine learning methods. (A) Dimensionality reduction using uniform manifold approximation and projection (UMAP) and Leiden clustering of CD4^+^ T cell subsets from all younger and older adults at the pre-vaccination timepoint. (B) Phenotype marker expression corresponding to each clustered population of CD4^+^ T cells. (C) Cell population frequencies were compared within or between age groups by Mann-Whitney U test with ns being not significant. (D) Identified regions of difference by age group with T-REX k-nearest-neighbor search and a 90% similarity threshold at the pre-vaccine timepoint. Significant pre-vaccine cell populations for older and younger adults are colored in blue and red, respectively. (E) T-REX cell frequency distribution by participant for significant populations. (F) CD4^+^ T helper populations identified by surface chemokine receptors CXCR3, CCR4, and CCR6 before vaccination compared between younger and older adults by Mann-Whitney U test. For statistical tests *p <0.05, **p <0.01, ***p <0.001, and ns being not significant.

Circulating T follicular helper (cTfh) cells are of particular interest because they aid B cell antibody responses upon influenza vaccination.^39,40^ We initially evaluated the relationship between humoral immune responses and the activation of cTfh cells defined as the expression of ICOS and CD38 (Supp. Fig. 2B) after vaccination. We found that changes in frequencies of circulating plasmablasts positively correlated with changes in activated cTfh cell frequencies in younger adults but not in older adults (Fig. 4A). Furthermore, activated cTfh cells, identified by ICOS^+^ CD38^+^, increased at days 7 and 28 post-vaccination in older adults, while younger adults showed no changes following vaccination (Fig. 4B). Frequencies of activated cTfh cells were significantly lower in older adults compared to younger adults pre-vaccination and post-vaccination (Fig. 4C). CXCR3^+^ Tfh cells, also called type 1 cTfh, have been shown to support the production of higher avidity antibodies to influenza.^40,41^ Type 1 cTfh cells were also present in our cohort at higher frequencies in younger adults, both before and after vaccination, compared to older adults (Fig. 4D). In summary, we demonstrate that there was no significant difference in major CD4^+^ T cell phenotypes between older and younger adults using unsupervised machine learning methods. However, subpopulations of T helper cells identified through biaxial gating differed between age groups, with older adult T helper cell phenotypes shifting away from antiviral T helper cells and with lower frequencies of cTfh populations compared to younger adults.

**Figure 4.**
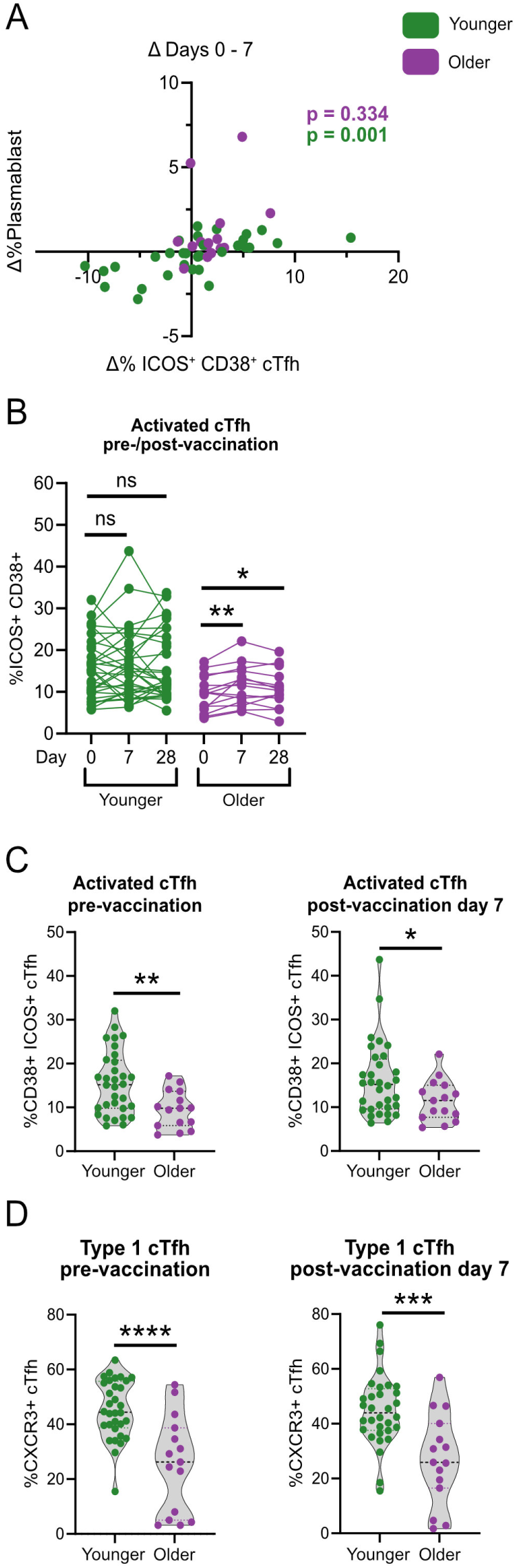
Older adults have decreased cTfh populations despite mounting a response post-vaccination. (A) Spearman correlation between the days 0-7 change in circulating plasmablasts and days 0-7 change in activated cTfh cells. Younger adult data points and statistics are colored in green, and older adults are colored in purple. (B) Changes in activated cTfh frequencies between days 0-7 and 0-28 by paired Wilcoxon rank test with Bonferroni correction. (C-D) Comparison of (C) activated cTfh and (D) type 1 cTfh cells by age group at day 0 and day 7 by Mann-Whitney U test. For statistical tests *p <0.05, **p <0.01, ***p <0.001, ****p <0.0001, and ns being not significant.

### Influenza-stimulated CD4^+^ T cells from older adults exhibit reduced transcriptional antiviral signature compared to younger adults

To better understand the differences between CD4^+^ T cell responses in older and younger adults, we repeated the AIM assay for 12 older and 12 younger adults and took a portion of the cells for single-cell RNA sequencing (scRNA-seq). Following integration, we identified major cell lineages such as B, T, natural killer, and myeloid cells. We then extracted, re-integrated, and re-clustered CD4^+^ T cells for improved resolution (Fig. S4A-B, Fig. 5A). We further classified CD4^+^ T cells based on well-characterized cell populations. These include naïve cells expressing high levels of *CCR7* and stimulation-induced naïve cells, termed interferon-stimulated gene (ISG) naïve cells due to their high expression of interferon-associated genes, such as IFIT1. Memory cells were classified either by major helper type (Th1, Th2, Th17, cTfh, and Treg) using hallmark genes such as *GATA3*, *FOXP3*, *IL7R*, and *CXCR5*, or into broad memory categories such as central memory (CM) or effector memory CD45RA^+^ (TEMRA), which are distinguished by lower *CCR7* expression. We also identified a unique cell population that appears primarily in stimulated samples, termed activated memory CD4 (act mem CD4), characterized by ISGs and *TNFRSF4* (encoding OX40) expression (Fig. 5A-B).

**Figure 5.**
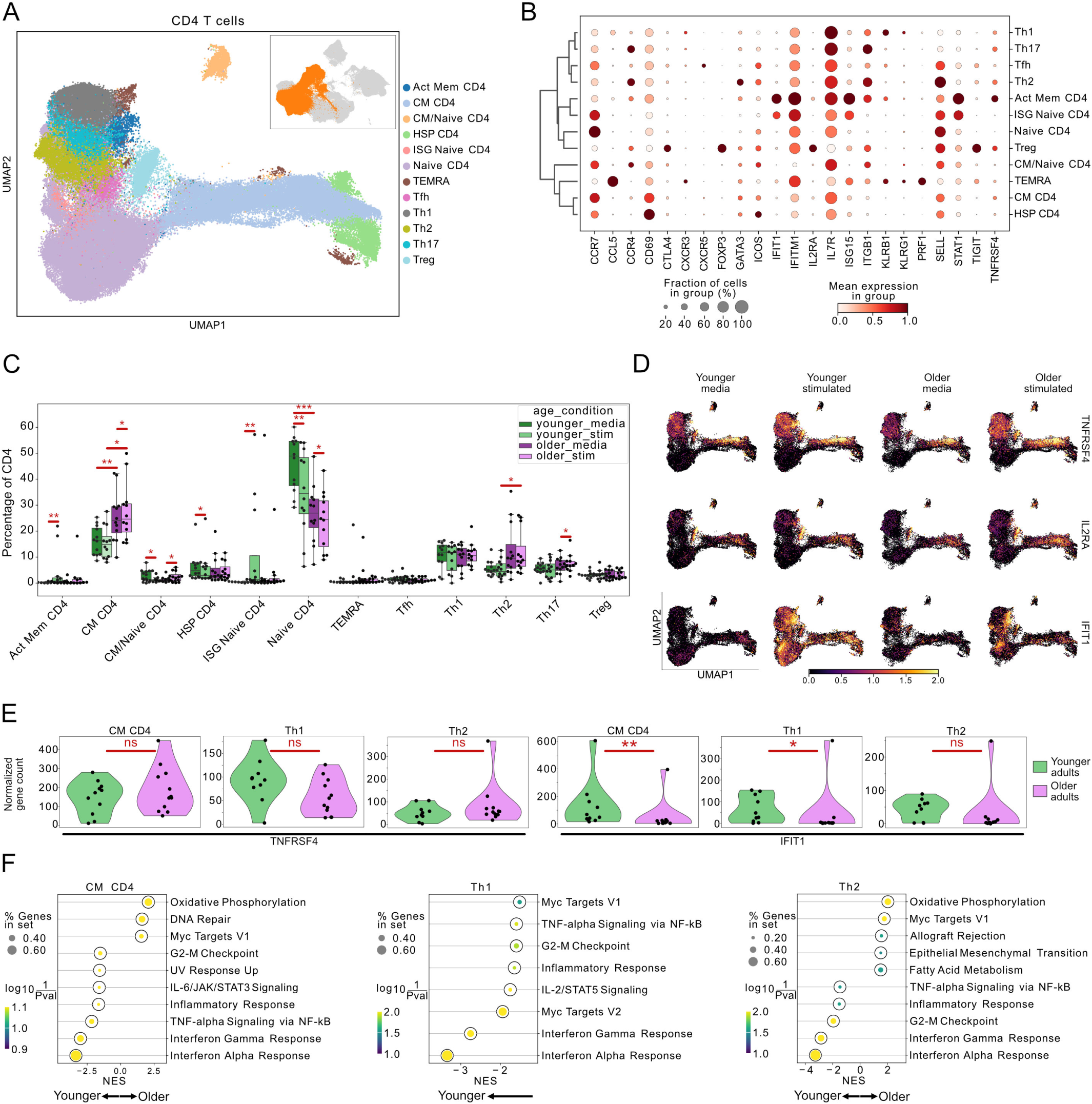
CD4^+^ T cells from older adults have altered compositional and transcriptional responses following stimulation. (A) UMAP of CD4^+^ T cells from all individuals with the inset UMAP exhibiting all PBMC with CD4^+^ T cells highlighted in orange. Cell population unabbreviated include act mem (activated memory), CM (central memory), HSP (heat-shock protein), ISG (interferon stimulated gene), TEMRA (T effector memory CD45RA^+^). (B) Relative mean gene expression dot plot of highlighted genes with higher expression indicated by dark red coloring, and cell fraction expression for genes indicated by circle size for CD4^+^ T cell populations. (C) Cell composition barplot split by age and stimulation condition and adjusted for individual contributions using a paired Wilcoxon rank or Mann-Whitney U test. (D) Gene expression projected on UMAP of TNFRSF4, IL2RA, and IFIT1 split between media-unstimulated and stimulated conditions. Bright yellow coloring indicates the highest expression, while black indicates no expression. (E) Violin plots for normalized TNFRSF4 or IFIT1 gene counts in stimulated memory T cell populations comparing older and younger adults with a Mann-Whitney U test. (F) Gene set enrichment dot plots using MsigDB Hallmark database comparing younger and older adult stimulated cell populations with a minimum FDR of 0.05. Gene sets in younger and older adults are negatively and positively enriched, respectively. For statistical tests *p <0.05, **p <0.01, ***p <0.001, ****p <0.0001.

To investigate changes during stimulation among older and younger adults, we employed a kNN search, comparing media and stimulated neighborhoods quantified by fold-change between the conditions. Both age groups had similar changes in cell populations; however, this method can be driven by outliers (Fig. S4C). We then compared cell type frequencies between age groups and in response to influenza antigen stimulation using a rank analysis to ascertain if changes were driven by select individuals. Younger adults had increased activated memory CD4 and ISG naïve CD4 T cells, while older adults had increased CM CD4, CM/naïve CD4, and Th17 T cells after stimulation. Both age groups showed decreased frequencies of naïve CD4 T cells after stimulation, with corresponding increases in ISG naïve CD4 and CM/naïve CD4 (Fig. 5C). Activated memory CD4 T cells are the primary antigen-reactive cell population by cell frequency, and these cells only change in response to antigen in younger adults with a mean of 0.322% (media) - 4.09% (stimulated). Although these cells are present in low numbers in older adults, they do not change in frequency with stimulation with a mean of 0.385% (media) - 2.09% (stimulated) (Fig. 5C & S4D). These cells are characterized by their low expression of *CCR7* and high expression of *TNFRSF4* and ISGs (Fig. 5B). Importantly, these influenza-reactive cells by scRNA-seq significantly correlated with the OX40^+^ CD40L^+^ influenza-reactive cells by flow cytometry, highlighting consistent findings across experimental methods (Fig. S4E).

Transcriptional changes occurred more broadly compared to compositional cell population changes after antigen exposure. CD4 T cell activation-induced genes, such as *TNFRSF4* and *IL2RA*, were broadly expressed across CD4^+^ cell populations when overlayed on the UMAP. Both age groups exhibited increased *TNFRSF4* transcript expression in helper and memory cells after antigen stimulation. *IFIT1*, known to be upregulated in T cells following T cell receptor stimulation^42,43^, was more highly expressed in CM CD4^+^ Th1 cells from younger adults after antigen stimulation (Fig. 5D-E).

To investigate broader changes in the transcriptome between younger and older adults, we employed pseudobulk differential gene expression coupled with gene set enrichment analysis (GSEA). We focused on three populations of memory T cells after antigen stimulation, including Th1, Th2, and CM cells. CM CD4 T cells from younger adults exhibited T cell activation-related gene set enrichment, such as interferon responses and TNF signaling. Older adult CM CD4 T cells had fewer enriched gene sets that met the false discovery rate (FDR) threshold, but included cellular processes for metabolism and proliferation. Th1 cells from older adults had no gene sets that met the FDR threshold. Th1 cells from younger adults displayed enrichment for cell proliferation, interferon responses, STAT5 signaling, and TNF signaling. Compared to younger adults, older adults had increased frequencies of CD4^+^ Th2 cells after antigen stimulation. Thus, we hypothesized that older adults responded to the antigen through Th2 cells instead of Th1. However, Th2 cells from older adults again displayed enrichment for gene sets involved in cell proliferation and metabolism. Th2 cells from younger adults also showed enrichment for G2/M cell cycle checkpoint but also displayed enrichment for interferon and proinflammatory responses (Fig. 5F). Overall, while both age groups displayed CD4^+^ T cell compositional changes with antigen stimulation, CD4^+^ T cells from older adults has less robust T cell activation-associated transcriptional patterns compared to younger adults.

### Myeloid cells from older adults exhibit reduced interferon signaling after stimulation

CD4^+^ T cells primarily rely on myeloid cells to interact with T cells through antigen presentation, resulting in T cell activation to perform effector functions. To better understand what is occurring in myeloid cells following antigen stimulation, we isolated myeloid cells from all PBMC in the scRNA-seq data object and reintegrated them to examine changes in finer detail (Fig. S5A-B). Within the myeloid compartment, we identified three main populations of monocytes, referred to as mono, and two types of dendritic cells (DC) (Fig. 6A). The monocyte populations include CD14 monocytes, CD16 monocytes, and interferon-stimulated gene (ISG) CD14 monocytes, characterized by the high relative expression of hallmark genes such as *FCGR3A* (encoding CD16), *CD14*, and interferon-induced genes like *STAT1* and *IFIT1*. Furthermore, we identified a small population of monocytes under stress or dying, showing significant upregulation of heat shock protein transcripts and DNA-damage-associated genes. Both DC populations exhibited elevated levels of antigen presentation and costimulatory genes, including *HLA-DR*, *CD40*, and *CD86*. Conventional DC (cDC) exhibited increased *CD40*, *CD86*, and *LYZ* expression. In contrast, plasmacytoid DC (pDC) showed high expression of interferon genes such as *IRF4*, *IRF7*, and *STAT1*, in addition to previously described pDC-specific genes like *GZMB* and *IL3RA* (Fig. 6B).^22,44,45^ To understand broad changes in these cell populations in response to antigen stimulation, we used a kNN search comparing neighborhoods found between media and stimulation. Myeloid cells from younger adults demonstrated an increased fold-change in the ISG CD14 monocyte region, whereas older adults showed an increased fold-change in the ISG CD14 monocyte and CD14 monocyte regions (Fig. 6C). Frequencies of these cell populations based on all myeloid cells were compared between age groups and stimulation conditions. Younger adults exhibited significant decreases in CD14 monocytes, increases in CD16 monocytes, and increases in ISG CD14 monocytes after antigen stimulation. Older adults, however, showed no changes in the frequency of these cells with stimulation but had a higher frequency of ISG CD14 monocytes compared to younger adults without stimulation and lower frequencies of cDC and pDC, both with and without stimulation, compared to younger adults (Fig. 6D).

**Figure 6.**
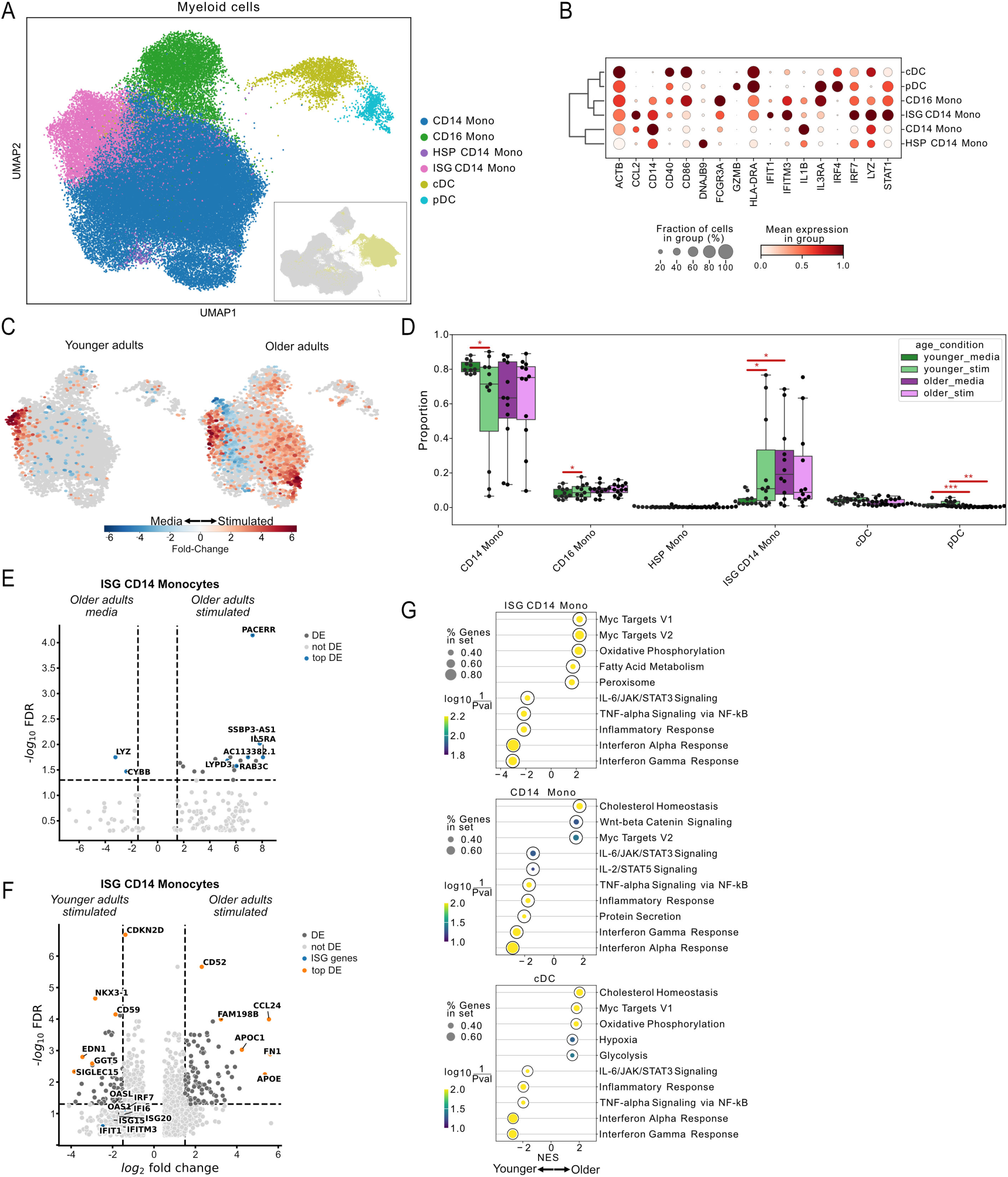
Older adults have reduced proinflammatory gene expression profiles in myeloid cell populations. (A) UMAP of myeloid cells from all individuals with the inset UMAP showing all PBMC with myeloid cells highlighted in yellow. (B) Relative mean gene expression dot plot for highlighted genes and their cell fraction expression for myeloid cell populations. (C) Differential abundance between media and stimulated cells was evaluated by kNN search in younger and older adults quantified by fold-change. (D) Cell composition barplot split by age and stimulation condition and adjusted for individual contributions using a paired Wilcoxon rank test when comparing media to stimulation or Mann-Whitney U test when comparing younger and older adults (*p <0.05, **p <0.01, ***p <0.001, ****p <0.0001). (E-F) Volcano plots of ISG CD14 monocytes with an FDR threshold of 0.05 and a minimum log_2_ fold-change of 1.5. (G) Gene set enrichment dot plots comparing stimulated cell populations from younger and older adults using the MsigDB Hallmark database with a minimum FDR of 0.05. Gene sets in younger and older adults are negatively and positively enriched, respectively.

Given that ISG CD14 monocytes were present in older but not younger adults prior to stimulation, we investigated how this population changes with age and response to antigen stimulation. By comparing media to stimulated ISG CD14 monocytes in older adults using pseudobulk differential gene expression, we found very few differentially expressed genes, with even fewer meeting significance criteria. Notably, no interferon genes were differentially expressed in either condition (Fig. 6E). Stimulated ISG CD14 monocytes exhibited significant differences in gene expression between older and younger adults. Younger adult cells expressed genes related to inflammation, proliferation, and interferon signaling, such as *GGT5*, *CDKN2D*, *EDN1*, and *OASL* (Fig. 6F). GSEA of stimulated ISG CD14 monocytes revealed significant enrichment of interferon responses, TNF signaling, and IL6-STAT3 signaling in younger adults. In contrast, older adult ISG CD14 monocytes exhibited no ISG genes and had higher expression of immunosuppressive genes such as APOE.^46^ ISG CD14 monocytes from older adults had significant enrichment of cellular processes related to proliferation and metabolism. This pattern was present in other cell populations that did not change in frequency with stimulation. CD14 monocytes and cDC exhibited similar enrichment patterns for interferon responses and other proinflammatory processes in younger adults, while older adults demonstrated gene expression patterns consistent with higher metabolic and proliferative activity (Fig. 6G). In summary, myeloid cells from older adults experienced less dynamic compositional changes after influenza antigen stimulation, and they lacked proinflammatory gene signatures consistent with antiviral responses compared to younger adults.

### Cell-to-cell communication inferencing reveals reduced transcriptional targets for immune-activating signals in older adults

We hypothesized that impaired signaling from antigen presenting cells might result in the reduced activation-associated transcriptional patterns observed in CD4^+^ T cells from older adults. We partitioned PBMCs by major cell types and merged the monocyte, dendritic, and CD4^+^ T cells into one data object to investigate cell-to-cell communication inferences based on gene expression. We used two different analyses: an intercellular communication method (Liana)^47^ to investigate ligand-receptor interaction events, where interactions were measured by the specificity and magnitude of the interaction across specific cells, and an intracellular method (NicheNet)^48^ to investigate downstream events after ligand-receptor interaction by ligand regulatory potential (Fig. 7A). The top 20 intercellular communication ligand-receptor pairs in younger adults show ligands for chemokines and cell-cell adhesion from myeloid cells. All myeloid cells express ligands for the galectin family in high magnitude and rank, which help modulate cell-cell adhesion. Monocyte-T helper cell interactions include *VCAN* and *CCL2*, whereas DC memory T cell interactions include *CCL22* and *CCL19* (Fig. 7B). Downstream effects of cell interactions were grouped with senders as myeloid cells and receivers as memory T cells. Ligands resulting in significant downstream transcriptional effects following receptor interaction are shown. Ligands on myeloid cells in younger adults signaling to T cells include cytokines such as *IL27*, *IL10*, *IL6*, *TNF*, and IL6 cytokine family members, including *CLCF1* and *OSM*. Other ligands produced fewer downstream effects, such as *SIGLEC15* and *CD48*. CD4^+^ T cells from younger adults had significant increases in interferon-induced genes (*OAS*, *IFIT*, and *TRIM* gene families), cell adhesion (*ICAM and CD82*), and immune response regulatory genes (*IKZF2* and *CCL4*) as a result of the myeloid cell interactions (Fig. 7C). Intercellular communication in older adults also included *CCL22*, *VCAN*, and galectin family ligands similar to younger adults. However, while older adults had increased HLA class II antigen expression, they had reduced chemokine signaling compared to younger adults (Fig. 7D). Myeloid cell ligands from older adults include *IL15*, *TGFB1*, and *TNF*. These ligands on myeloid cells induced cytokine expression and cell activation through receptors, such as *IL1A* and *NFKB2*, on CD4^+^ T cells from older adults (Fig. 7E). While both age groups exhibited cell-cell adhesion interactions, younger adults had increased chemokine interaction between monocyte-CD4^+^ T cell populations, whereas older adults had pathways suggesting increased antigen presentation to T cells. Despite the increased antigen presentation, older adults had reduced downstream transcriptional effects and interferon signaling.

**Figure 7.**
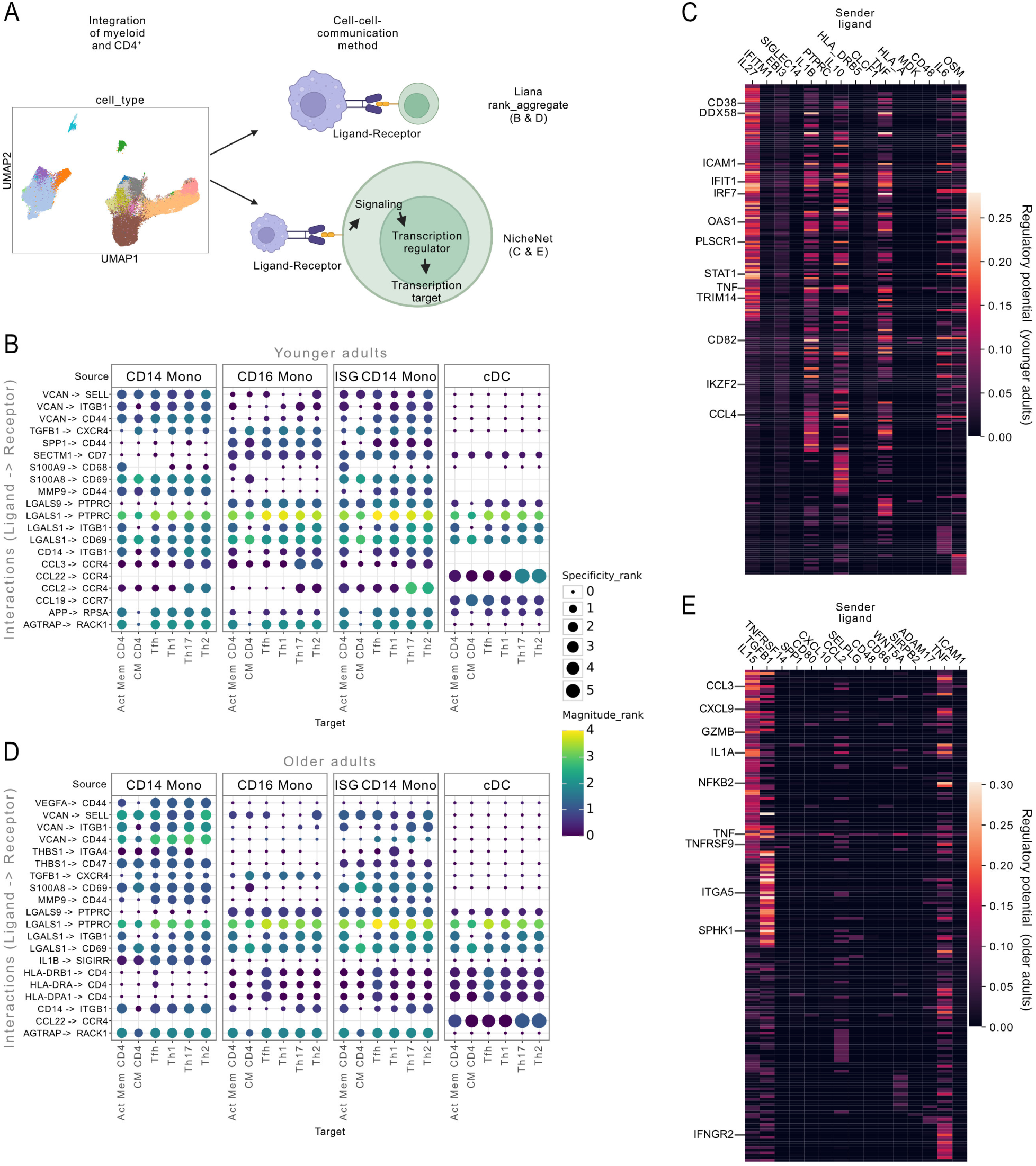
Older adults have altered intercellular communication programming between CD4^+^ T cells and myeloid cells compared to younger adults. (A) Representative diagram workflow following integration for inter and intracellular communication inferencing. (B, D) Liana cell-cell-communication inferencing within stimulated cells for younger adults (B) or older adults (D). The top 20 significant ligand-receptor pairs are shown from Liana, with myeloid cells being the sender (source) and memory CD4^+^ T cells being the receiver (target). Only statistically significant interactions are shown. (C, E) NicheNet communication inferencing comparing media to stimulated cells showing top interactions in stimulated cells in (D) younger adults and (E) older adults. Sender and receiver cell populations were grouped. The top ligands on the x axis from myeloid cells (sender) result in downstream signaling and transcriptional targets on the y axis in memory CD4^+^ T cells (receiver), measured by ligand regulatory potential. Downstream gene targets were annotated based on representatives from gene families and relevance to immune responses such as cytokines, chemokines, immune regulation, and antiviral activity.

## DISCUSSION

Despite the administration of improved vaccine formulations, older adults are still at a disproportionate risk of influenza-related complications compared with younger adults. While these vaccines elicit comparable antibody responses between age groups (as shown in Fig. 1) it is clear that this alone is not sufficient to prevent disease in older adults. Despite the fact that antigen-specific CD4^+^ T cell responses have been correlated with improved protection and decreased disease severity in humans, antigen-specific T cell responses following vaccination are significantly less studied, especially in at-risk populations. Therefore, it is crucial that we broaden our understanding of vaccine-induced T cell immunity.^49^ We investigated factors that underlie differential innate and adaptive responses to antigen stimulation between older and younger adults using scRNA-seq and cytometric surface marker characterization before and after vaccination. We found that both age groups mounted similar HA-antibody responses, but older adults had reduced CD4^+^ T helper responses after influenza antigen stimulation compared to younger adults. These reduced cellular responses were not associated with well-described features of an aging immune system, such as the frequency of senescent CD4^+^ T cells or CMV seropositivity. Instead, we observed reduced frequencies of influenza reactive CD4^+^ T cells and attenuated transcriptional interferon induction following stimulation in myeloid and CD4^+^ T cells in older adults. This suggests that myeloid cells are a major contributor to reduced cellular responses observed with aging.

As shown in Fig. 1, antibody responses to vaccine antigens were comparable between age groups, likely because older adults received improved formulations designed to boost HA antibody responses.^5,6,16^ Although the current influenza vaccines are designed to elicit antibody responses and not CD4^+^ T cell responses, improved formulations, such as MF-59 adjuvanted vaccines, for older adults have been shown to induce antigen-specific T cell responses in older adults.^25^ Despite our cohort of older adults receiving different versions of vaccines with reported superior immunogenicity, they still developed poor cellular responses compared to younger adults who received the standard dose vaccine.

Broad changes in T cell phenotypes, notably cTfh responses, are associated with aging and impaired vaccine-specific immune response.^17,30,50^ Using supervised and unsupervised analysis techniques, we found no significant differences in large cell populations, including memory, naïve, and senescent cells. T helper subsets are integral in the coordination of both myeloid and CD8^+^ T cells during an immune response. T helper subsets shifted away from an antiviral phenotype in older adults. Many current vaccines, including influenza, primarily use aluminum salts as an adjuvant, inducing a type 2 immune response.^51–53^ Since older adults have likely been vaccinated for several years, this could explain the observed shift toward a Th2 phenotype upon recall of these responses due to repeated vaccinations or pathogen exposure. We also observed reduced cTfh cell population frequencies in older adults compared to younger adults. Activated cTfh cells have been previously correlated with B cell and serologic antibody responses following influenza vaccination in younger and older adults.^26,29,30,39,54^ We did not observe this correlation in older adults; however, this could be due to our limited sample size.

Previous research examining transcriptional differences in T cell responses to influenza vaccination in older adults has focused on cTfh cells *ex vivo*, revealing increased TNF inflammatory transcriptional patterns compared to younger adults.^29,30^ While these studies found increased TNF transcriptional signatures in this specific T cell subset in older adults post-vaccination, we show that reduced interferon transcriptional signatures are broadly conserved among T cells from older adults after stimulation. This observation likely supports the inflammaging theory, where basal-level inflammation could cause impaired immune responses.

While cTfh cells are important for specific vaccine responses, the broader memory CD4^+^ T cell population plays a crucial role in immunity and pathogen clearance. More recently, studies have highlighted the importance of memory CD4^+^ T cell responses and interferon induction in myeloid and lymphoid cells for protection from influenza-related disease in a clinical setting.^55,56^ Moreover, immune responses to both SARS-CoV-2 infection and vaccination have shown reduced T cell activation and interferon responses in older adults, which is consistent with our findings.^57,58^ Both our flow cytometric results and single-cell transcriptional analysis of CD4^+^ T cells revealed that older adults produce fewer influenza-specific CD4^+^ T cells after vaccination than younger adults. We also observed reduced ISG induction across many CD4^+^ T cell subsets in older adults compared to younger adults. Prior work found a reduction of type I interferon signaling in naive CD4^+^ T cells from older adults after direct TCR stimulation with toxic shock syndrome toxin.^42^ Our work significantly expands this work to encompass naïve and memory CD4^+^ T cells in the context of influenza.

Other groups have found that elevated basal levels of proinflammatory signatures in myeloid cells negatively affect vaccine responses due to their reduced ability to further induce inflammation for an immune response.^59–61^ Monocytes from older adults have been described to have reduced ISGs such as TRAF3, which in turn reduces RIG-I signaling, which is important for an antiviral response.^62^ In our work, transcriptional analysis of myeloid cells revealed a stimulation-specific ISG monocyte population in younger adults, present with and without stimulation in older adults. Despite clustering with monocytes highly expressing ISGs from younger adults, these ISG CD14 monocytes from older adults likely represent a chronic proinflammatory monocyte subset.

Cell-cell interactions between antigen presenting cells, CD4^+^ T cells, and B cells are critical for coordinating effective adaptive immune responses for initial pathogen recognition and long-lived immunity. Our CCC inferencing analysis between myeloid (sender) and memory T (receiver) cells revealed significant differences in chemokine interactions and downstream proinflammatory signaling between younger and older adults. Notably, the primary cytokine ligand from the myeloid cells (IL27 and IL6 in younger adults vs IL15 in older adults) highlights the programming differences in myeloid cells between the age groups. Although this is an inference analysis and needs to be experimentally validated, the significant reduction in interferon signaling was consistent with the reduced CD4^+^ T cell transcriptional signature of interferon activation in older adults.

In this study, we found that older adults had reduced frequencies of influenza-reactive CD4^+^ T cells and attenuated interferon induction after stimulation with influenza antigen in myeloid and CD4^+^ T cells after vaccination. Our study suggests that myeloid cells are a major contributor to diminished CD4^+^ T cell responses observed with aging, and formulating vaccines to induce tailored myeloid responses might lead to improved vaccine responses and outcomes in older adults. Further investigation into the induction of myeloid cell activation and presentation to antigen-specific CD4^+^ T helper responses in at-risk populations is necessary for future influenza vaccine improvements. Transcriptional signatures defined in this study could potentially identify individuals at an increased risk for infection and further infection-related medical complications.

## STUDY LIMITATIONS

The detailed single-cell profiling we performed limited our sample size, so it was not possible to control for all study population differences. We controlled for sex assigned at birth (referred to as ‘sex’ in this study) by having equal proportions in each age group; however, racial and ethnic demographics could not be controlled because of the modest cohort size and limited statistical power. Although all participants received an influenza vaccine in the prior year, we cannot determine how many years prior individuals received influenza vaccines. Receiving an influenza vaccine the prior year was not an inclusion/exclusion criterion. Furthermore, we could not compare results from older adults vaccinated with a standard-dose vaccine, as the formulations given in this cohort are the current standard of care.

Antibody titers to influenza viruses reported in this study used HAI. Thus, we cannot discern non-sialic acid binding site antibody responses between age groups. Despite the fact that the older adults in our cohort are over 65, they likely represent relatively healthy individuals. Further study of differences in immune senescence in older adults with frailty and significant comorbidities is needed. Additionally, we further reduced our cohort size for scRNA-seq due to assay limitations. From the scRNA-seq data, we collected over 500,000 cells, but some annotated cell populations had low counts when separated by the individual; because of this, gene expression analysis was not possible. Future studies should focus on expanding cohort demographics and investigating the effect of new vaccine platforms on CD4^+^ T cell and myeloid cell responses.

## RESOURCE AVAILABILITY

### Lead contact

Further information and requests for resources and reagents should be directed to and will be fulfilled by the lead contact, Spyros A. Kalams.

### Materials availability

This study did not generate new or unique reagents.

### Data and code availability

No original code is reported in this work. Additional information or code to aid in reanalyzing the data reported in this work is available upon request to the lead contact, Spyros A. Kalams.

## ACKNOWLEDGMENTS

We thank the individuals for participating in this study. We thank Rita M. Smith for repository management; Christian M. Warren for flow cytometry guidance; Eric Alves, Silvana Gaudieri, Mona Mashayekhi, Celestine N. Wanjalla, Ty Sornberger, and Chelsea N. Campbell for comments on the manuscript. Flow cytometry data were collected in the VUMC Flow Cytometry Shared Resource core facility, supported by the Vanderbilt-Ingram Cancer Center (P30 CA68485) and the Vanderbilt Digestive Disease Research Center (DK058404). Mass cytometry data were collected in the Vanderbilt Mass Cytometry Center of Excellence, supported by Vanderbilt-Ingram Cancer Center (P30 CA068485). Single-cell sequencing data were collected in partnership with Vanderbilt Technologies for Advanced Genomics. VANTAGE is supported by CTSA grant 5UL1 RR024975-03, the Vanderbilt-Ingram Cancer Center (P30 CA068485), the Vanderbilt Vision Center (P30 EY08126), and NIH/NCRR (G20 RR030956). For work described in this manuscript, S.A.K., J.M.O. were supported by the National Institute of Allergy and Infectious Diseases award R01 AI142095.

## AUTHOR CONTRIBUTIONS

J.M.O. and S.A.K. developed the methodology. H.K.T., J.L.C., N.B.H., and S.A.K. designed the clinical study, and J.E. performed sample collection. J.M.O., J.D.S., C.H-N., and L.K. performed experimentation and data analysis. J.M.O. wrote the original draft. J.D.S, C.H-N., J.L.C., S.S.B, and S.A.K. reviewed and edited the manuscript. S.A.K. acquired funding and oversaw the work.

## Materials and Methods

### Cohort description

To assess immune responses in older adults, we recruited 15 older adults and 32 younger adults during the 2019-2020 Northern Hemisphere influenza season. Whole blood specimens were collected on the day of vaccination (day 0) and post-vaccination on days 7 and 28 during the fall of 2019, with peripheral blood mononuclear cell isolation immediately following each collection. Younger and older adults were similar in terms of the percentage of females (81% and 60%) and those identifying as white (81% and 93%), but differed by median age (32.7 and 70.3 (Table S1). Individuals younger than 45 received a quadrivalent Fluzone standard-dose vaccine containing H1N1, H3N2, B-Victoria, and B-Yamagata, while adults ≥65 received either a trivalent Fluzone high-dose, FludAD, or Flublok vaccine which did not contain B-Yamagata. We measured CMV antibody titers by ELISA in older and younger adults (Fig. S3A). Older adults had a higher rate of CMV seropositivity (73%) compared to younger adults (44%) Fig. S3B). All individuals provided written informed consent.

### Blood sample collection and processing

This study was approved by the Vanderbilt University Medical Center Institutional Review Board and all donors by written informed consent (IRB 161647 and 191639). Blood was drawn in ETDA coated tubes (Becton Dickinson) at pre-vaccination (day 0), 7, and 28 days post-vaccination in the fall of 2019. All blood processing was performed at room temperature (20°C-25°C). Whole blood was centrifuged at 400 x g for 10 minutes without brake. Plasma was collected and further centrifuged at 1200 x g for 10 minutes with brake and then aliquoted and frozen at -80°C. The plasma pellet formed was taken for downstream HLA typing. Phosphate buffered saline (PBS) was added to whole blood, returning the contents to its original volume. Blood was transferred to 50mL Leukosep tubes with Ficoll Paque andcentrifuged at 800 x g for 15 minutes with no brake. The PBS-buffy coat was collected and added into a new 50mL non-barrier conical containing 45mL of PBS followed by centrifugation at 400 x g for 10 minutes. The supernatant was removed without disturbing the cell pellet. Cell pellets were resuspended in a small volume to combine pellets from multiple tubes. PBS was added to maximum volume in a conical tube and centrifuged at 400 x g for 10 minutes. The supernatant was aspirated, and the cells were resuspended at 10 million cells per milliliter in cold cryoprotective solution (10% DMSO, 90% FBS) for overnight freezing at -80°C. Frozen cells were transferred from the -80°C freezer to liquid nitrogen the following day. Concurrently, blood was collected in an SST tube (Becton Dickinson) and allowed to clot for 30 minutes. SST tubes were spun at 1100 x g for 10 minutes. Serum was collected, aliquoted, and frozen at -80°C.

### Hemagglutination Inhibition (HAI) assay

Serum was treated with receptor destroying enzyme (Denka Seiken) for 18 hours at 37°C followed by inactivation at 56°C for 45 minutes. Serum was diluted with PBS to a final dilution factor of 1:10. Turkey erythrocytes (Lampire Biologics) were prepared by washing 2 times with PBS and resuspended in a final concentration of 0.5%. For serum hemadsorption, 100μL of 0.5% washed turkey erythrocytes were centrifuged at 360 x g for 5 minutes, and the supernatant was aspirated. Diluted serum was combined with pelleted erythrocytes and incubated for 1 hour at 4°C while inverting every 15 minutes to mix. Serum was centrifuged, and the supernatant was saved. Antigens (A/Brisbane/02/2018, A/Kansas/14/2017, B/Colorado/06/2017, B/Phuket/3073/2013) were titrated for 4 HA units/25uL confirmed with a back titration in a V-bottom plate. The treated serum was serially diluted 2-fold in duplicates, followed by the addition of an equal volume of 4 HA/25uL antigen, totaling 50uL. V-bottom plates were incubated for 30 minutes at room temperature. 50uL of 0.5% turkey erythrocytes were added and incubated for 30-45 minutes at room temperature. Antibody titer was read as the last well exhibiting hemagglutination. If duplicates showed a greater than 2-fold variation, the assay was repeated.

### Activation Induced Marker (AIM) assay

Peripheral blood mononuclear cells (PBMC) were thawed in 9mL of PBS and 15μg of nuclease S7. After 2 washes in 10mL of PBS centrifuging at 420 x g, cells were rested for 4 hours in RPMI containing 10% human serum, 1X penicillin streptomycin, 10mM HEPES buffer, and 2mM L-glutamine at 37°C in 5% CO_2_. Subsequently, cells were incubated with 0.5ug/mL CD40 blocking antibody for 15 minutes at 37°C. 1.5-2 million blocked cells were added to each stimulation condition and incubated with media or stimulant and 0.03ug/mL of CD40L-PE antibody in a 96-well U-bottom plate. Stimulants include inactivated virus A/Brisbane/02/2018 and A/Kansas/14/2017 normalized to 16 HA units. Cells were incubated for 18 hours at 37°C with 5% CO_2_. Cells were washed and resuspended in PBS for flow cytometry staining. A master mix of fluorochrome conjugated antibodies (Table S2) was added to each condition and incubated for 20 minutes at room temperature, followed by 1 wash in PBS and fixation using BD stabilizing fixative. Samples were acquired on a BD LSR Fortessa. Data events were gated by time to ensure quality data events for downstream analysis. CD4^+^ T cells were identified by live/dead aqua^−^, CD3^+^, and CD19^−^. Memory cells were identified by either CD45RO^+^ CCR7^+/-^, or CD45RO^−^ CCR7^−^. Influenza-specific cells were identified by OX40^+^ CD40L^+^ dual positivity. An example gating strategy can be viewed in Fig. S2A.

### Mass cytometry data acquisition

PBMC were thawed in PBS and 15ug of S7 nuclease, followed by 2 washes in PBS, centrifuging at 420 x g. Cells were resuspended in PBS and stained with 2μL/mL Rh-103 viability at 37°C for 15 minutes. Cells were washed twice with PBS containing 1% BSA and resuspended in PBS with 1% BSA. Antibody master mix (Table S2) was added to each sample and incubated at room temperature for 30 minutes with a total volume of 100μL. Cells were subsequently washed twice with PBS and fixed with 1.6% PFA final concentration for 10 minutes, followed by washing with PBS and resuspended in 1mL cold methanol and stored at - 20°C. The day before running on the Fluidigm Helios, cells were washed with PBS and resuspended with PBS/1% BSA. Cells were intracellularly stained for Ki-67 for 20 minutes at room temperature. 2μL of 25μM DNA intercalator (Ir) and a final concentration of 1.6% PFA was added to cells and incubated for 20 minutes at room temperature, then placed at 4°C. Before running on the instrument, samples were washed with PBS, centrifuged, and resuspended in H_2_O. EQ four element calibration beads (Standard Biotools) were added to each sample.

### Mass cytometry data analysis

Signal normalization across the file set using 5-channel positive control beads (premessa R package). Data was manually gated for CD45+ live singlet cells as previously outlined.^63^ These events are non-bead, viable, low to medium event length, and have a Gaussian pulse. CD45+ events were gated against time to avoid minor clog events and compromised data. Quality live cell events were gated on CD3^+/-^ and CD19^+/-^ to avoid events beyond axis limits. Events were downsampled for each sample to approximately 45,000 quality data events. Data files were arcsine transformed and concatenated using the flowcore R package. CD4^+^ T cells were selected for Uniform Manifold Approximation and Projection (UMAP) dimension reduction with a k-nearest-neighbor (kNN) of 40 using and mininmum distance of 0.25 followed by kNN search and Leiden algorithm clustering with a resolution of 0.7.^64,65^ Cluster population percentages were exported, and plots were made using ggplot2. T-REX was employed to identify differences in cell populations between age groups by creating a UMAP followed by a kNN search between two conditions. Hotspots are identified by a set threshold (90% in this study) unique to one sample.^38^ All analyses were performed using R 4.3.2.

### CMV ELISA

We followed the protocol provided by AVIVA Systems Biology GWB-892399. Briefly, serum samples from pre-vaccination (day 0) were heat inactivated at 56°C for 45 minutes. Inactivated serum was diluted at 1:20, 1:40, or 1:100 and incubated in pre-coated wells with CMV purified protein. Serum was incubated in duplicates at 37°C for 30 minutes, followed by washing 5 times with 0.05% PBS-tween. Concurrently, CMV IgG standards were also incubated at 37°C in duplicates. Following washing, samples and standards were incubated with anti-IgG conjugates HRP antibody. Samples and standards were washed again 5 times with 0.05% PBS-tween, followed by the addition of TMB reagent and incubation at 37°C for 15 minutes. An equal volume of stop solution (1N HCl) was added, and absorbance was read at 450nm within 15 minutes.

### HLA and KIR typing

DNA from plasma pellets was extracted using a Qiagen DNeasy Blood and Tissue kit. Extracted DNA was sent to Immunogenomics, Immunology, and Infectious Disease lab at Murdoch University (Perth, WA, Australia) for KIR and HLA typing. Primers designed to target locus specific exons were pooled in a multiplexed PCR assay. Amplicons were combined in equivalent concentrations for balanced read coverage. Sequencing was performed on an Illumina MiSeq using a paired-end 300bp kit (Illumina, CA, USA). Reads were filtered and demultiplexed, followed by alignment to a reference sequence containing all KIR genes with CLCbio genomics workbench (QIAGEN Bioinformatics) or a proprietary allele calling algorithm. using the latest IMGT HLA allele database. The current IPD-KIR or IMGT HLA allele database was used for KIR or HLA allele identification, respectively.^66,67^

### Single-cell RNA sequencing

PBMC were thawed and stimulated as outlined above in an AIM assay. Cells were counted and pooled to include two individuals per sample in equivalent portions before droplet and library generation using the 10X Genomics Next GEM 5’ V2 kit per the manufacturer’s instructions. We pooled an unstimulated (media) and stimulated sample to increase heterogeneity and reduce batch effects. After library generation, sequencing was performed on a NovaSeq 6000 S4 paired-end 150bp flow cell targeting 40,000 reads per cell.

### Bioinformatic processing of single-cell RNA sequencing data

CellRanger version v7.1.0 or v7.2.0 was used to align reads to a KIR-modified GRCh38, correcting for KIR misalignment.^68^ There were no significant updates with CellRanger related to gene expression that would impact our data quality. Cells containing <500 total genes, <200 unique genes, or more than 10% mitochondrial genes were eliminated. Samples were demultiplexed using Souporcell, and intersubject doublets were identified.^69^ Samples were matched to the corresponding participant’s HLA genotype by arcasHLA pairing. Cells not assigned “singlet” by Souporcell were dropped. Raw counts were adjusted using SoupX and then rounded to integers. Further doublet discrimination was performed using scDblFinder to identify intra-sample doublets. Cells with scores above the fit minimum were dropped. Data were integrated using the scVI method with ray tune hyperparameter tuning. Integer-rounded SoupX counts were used for highly variable feature selection and integration model training. The batch key was set to multiplexed sequencing library prep with participant ID as a categorical covariate, followed by percent counts from MT genes and total counts were used as continuous covariates for scVI model training. The integration model latent space was used for Leiden clustering at resolution three and UMAP visualization. Leiden clusters were assigned to one of three major lineage groups based on top gene markers: T/NK cell, B cell, or Myeloid cell. Each major cell population was subsetted, reintegrated, and re-clustered for improved granularity. After integration, cell clusters were annotated using CellTypist and gene expression using rank_genes_groups from scanpy. Composition analysis was done using MiloR implementation in pertpy with default parameters^70^ or by paired Wilcoxon rank test from cell frequencies. MiloR kNN search was set to 30, and use_rep was scVI. The neighborhood plot alpha value was set to 0.1, and the minimum size was set to 5. Scanpy dotplot was used for relative expression on normalized counts with a predefined gene list and the standard_scale set to var. Violin plots were created using the scanpy plot violin function using normalized counts from pseudobulk cell populations by individual, followed by a Mann-Whitney U test in numpy. We subset each cell population and then pseudobulk group by individual with a minimum cell count of 30. We required 75% of individuals within the comparator groups to have a pseudobulk data point for any given cell population before performing downstream analysis. Differential gene expression was performed using pyDEseq2 followed by GSEA analysis with gseapy. Volcano plots of differentially expressed genes were used with the Sanomics Python package. The GSEA prerank method was used in combination with the MSigDB_Hallmark_2020 database in conjunction with the GO_Biological_Process_2023 for verified findings. Cell-cell communication (CCC) analysis was performed using Liana and NicheNet. Liana can only infer CCC in a single group, so the data was subset by age and condition, followed by pseudobulking by cell type. We used the rank_aggregate method on log normalized counts and a filtered_lambda threshold of 0.05. Source labels included CD14 Mono, ISG CD14 Mono, CD16 Mono, and cDC. Plasmacytoid DCs were excluded due to low cell numbers in some older adults. Target labels included CM CD4, Th1, Th2, Th17, and Tfh cells. Activated memory CD4 cells were also excluded, as older adults had low cell numbers. We then compared media to stimulated CCC within each age group using NicheNet. We used https://zenodo.org/record/7074291/files/ligand_target_matrix_nsga2r_final.rds for the ligand target matrix and https://zenodo.org/record/7074291/files/lr_network_human_21122021.rds for the ligand-receptor network reference. We designated the sender and receiver cells in the same way we did with Liana. Cell populations were pseudobulked by individuals prior to running the inferencing analysis. The threshold for the p-value was set to 0.05, and the log fold-change was set to 1. Annotated ligand target genes are representative of overall changes following CCC inferencing.

