## Supplemental Table 1 for "Antiviral CD4^+^ T and myeloid cell responses to influenza vaccines are attenuated in older adults"

Older adults must have been 65 or older, while younger adults must have been 45 or younger upon enrollment. Younger adults were selected at random from a large cohort of vaccinated individuals.

| | Older Adults | Younger Adults |
|---|---|---|
| Number of Participants | 15 | 32 |
| Age Range | 65 - 77.7 years | 19.3 - 44.6 years |
| Median Age | 70.3 | 32.7 |
| Sex (Female/Male) | 9/6 | 20/12 |
| Black | 0 | 4 |
| Asian | 0 | 1 |
| White | 14 | 26 |
| Mixed/Other | 1 | 1 |
| Hispanic | 1 | 2 |
