## Supplemental Table 2 for "Antiviral CD4^+^ T and myeloid cell responses to influenza vaccines are attenuated in older adults"

Supp Table 1. Cytometry antibodies used

Fluorescent flow cytometry

| Target | Fluorphore | Clone | Final Concentration |
|---|---|---|---|
| PD1 | APC | EH12.2H7 | 1.0µg/mL |
| CD3 | AlexaFluor 700 | UCHT1 | 0.5µg/mL |
| CD45RO | PE CF594 | UCHL1 | 0.2µg/mL |
| OX40 | PE-Cy7 | BER-ACT35 | 0.2µg/mL |
| CD40L | PE | 24-31 | 0.03µg/mL |
| CD4 | PerCP-Cy5.5 | RPA-T4 | 1.0µg/mL |
| CXCR5 | AlexaFluor 488 | RF8B2 | 1.3µg/mL |
| CCR7 | Brilliant Violet 421 | 150503 | 1.5uL |
| CD25 | Brilliant Violet 786 | M-A251 | 0.25µg/mL |
| CD86 | Brilliant Ultraviolet 737 | FUN-1 | 0.2µg/mL |
| CD19 | Brilliant Ultraviolet 395 | SJ25C1 | 0.1µg/mL |
| Viability | Live/Dead Aqua | NA | 1:1000 dilution factor |

Mass cytometry

| Target | Metal Tag | Tag Isotope | Volume to add (ul) | Standar biotools catalogue number |
|---|---|---|---|---|
| CD45 | Yttrium (Y) | 89 | 1.0µL | 3089003B |
| Live/dead | Rhodium (Rh) | 103 | 2.0µL | 3103 |
| CCR6 | Praseodymium (Pr) | 141 | 0.5µL | 3141003A |
| CD38 | Neodymium (Nd) | 144 | 0.5µL | 3144014B |
| CD4 | Neodymium (Nd) | 145 | 1.0µL | 3145001B |
| CD8a | Neodymium (Nd) | 146 | 0.1.0 | 3146001B |
| CD14 | Neodymium (Nd) | 148 | 1.0µL | 3148010B |
| CD45RO | Samarium (Sm) | 149 | 0.5µL | 3149001B |
| CD86 | Neodymium (Nd) | 150 | 0.5µL | 3150020B |
| ICOS | Europium (Eu) | 151 | 0.5µL | 3151020B |
| CD21 | Samarium (Sm) | 152 | 1.0µL | 3152010B |
| CCR2 | Europium (Eu) | 153 | 0.5µL | 3153023B |
| TIGIT | Samarium(Sm) | 154 | 0.5µL | 3154016B |
| PD-1 | Gadolinium (Gd) | 155 | 2.0µL | 3155009B |
| CXCR3 | Gadolinium (Gd) | 156 | 0.5µL | 3156004B |
| OX40 | Gadolinium (Gd) | 158 | 0.5µL | 3158012B |
| CCR7 | Terbium (Tb) | 159 | 0.25µL | 3159003A |
| CD28 | Gadolinium (Gd) | 160 | 1.0µL | 3160003B |
| Ki-67 | Dysprosium (Dy) | 161 | 1.0µL | 3162007B |
| CD69 | Dysprosium (Dy) | 162 | 0.1µL | 3162001B |
| BTLA | Dysprosium (Dy) | 163 | 0.5µL | 3163009B |
| CXCR5 | Dysprosium (Dy) | 164 | 0.5µL | 3164029B |
| CD19 | Holmium(Ho) | 165 | 0.5µL | 3165025B |
| CD27 | Erbium (Er) | 167 | 0.5µL | 3167006B |
| CD138 | Erbium (Er) | 168 | 1.0µL | 3168009B |

| | | | | |
|---|---|---|---|---|
| CD25 | Thulium (TM) | 169 | 0.25µL | 3169003B |
| CD3 | Erbium (Er) | 170 | 2.0µL | 3170001B |
| CD20 | Ytterbium (Yb) | 171 | 0.5µL | 3171012B |
| CD57 | Ytterbium (Yb) | 172 | 0.5µL | 3172009B |
| CD137 | Ytterbium (Yb) | 173 | 1.0µL | 3173015B |
| HLA-Dr | Lutetium (Lu) | 174 | 1.0µL | 3174001B |
| CCR4 | Lutetium (Lu) | 175 | 0.5µL | 3175035A |
| CD127 | Ytterbium (Yb) | 176 | 2.0µL | 3176004B |
| Nuc acid | Iridium (Ir) | 191/193 | 2.0µL | 201192A |
| CD16 | Bismuth (Bi) | 209 | 0.5µL | 3209002B |
