## Supplementary figures and images for "Antiviral CD4^+^ T and myeloid cell responses to influenza vaccines are attenuated in older adults"

### Supplementary Figure 1

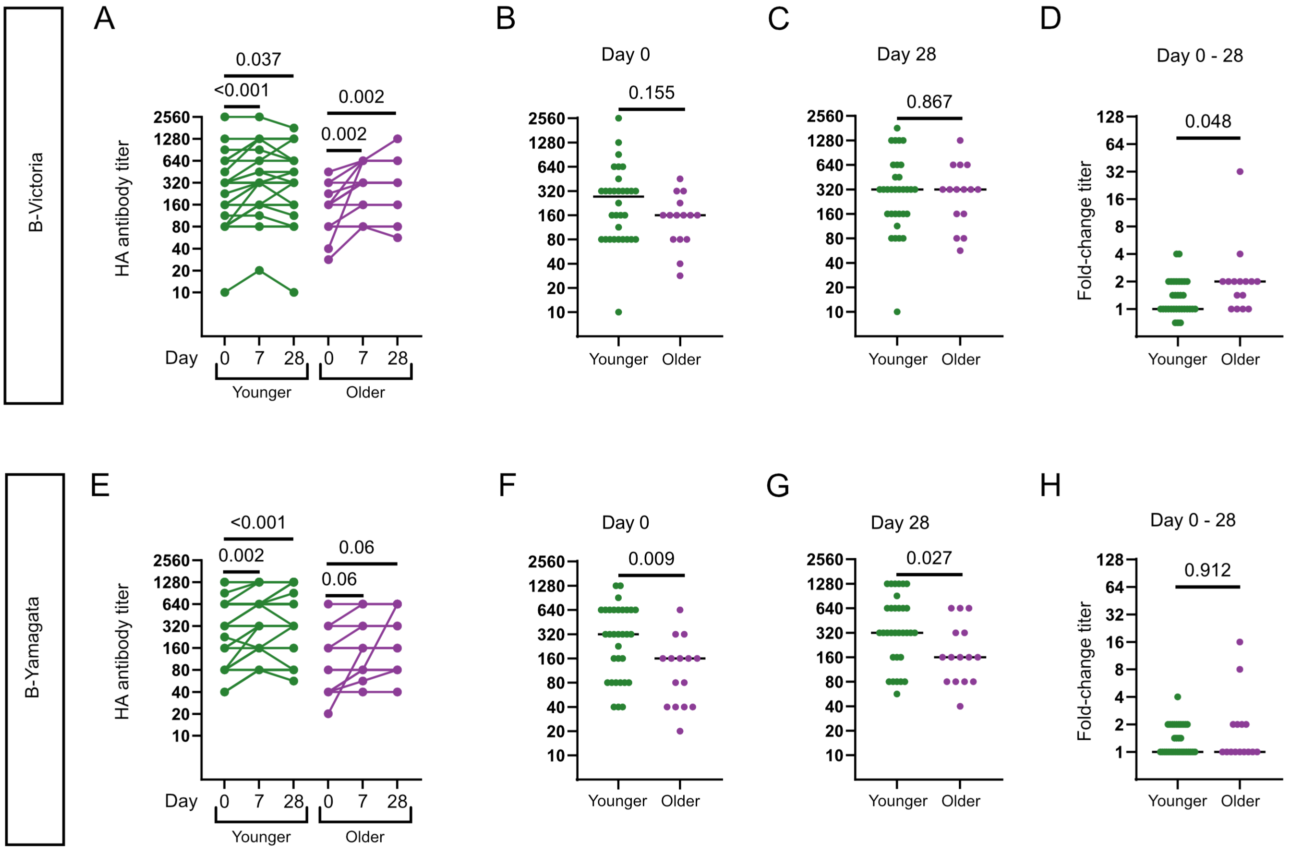

### Supplementary Figure 2

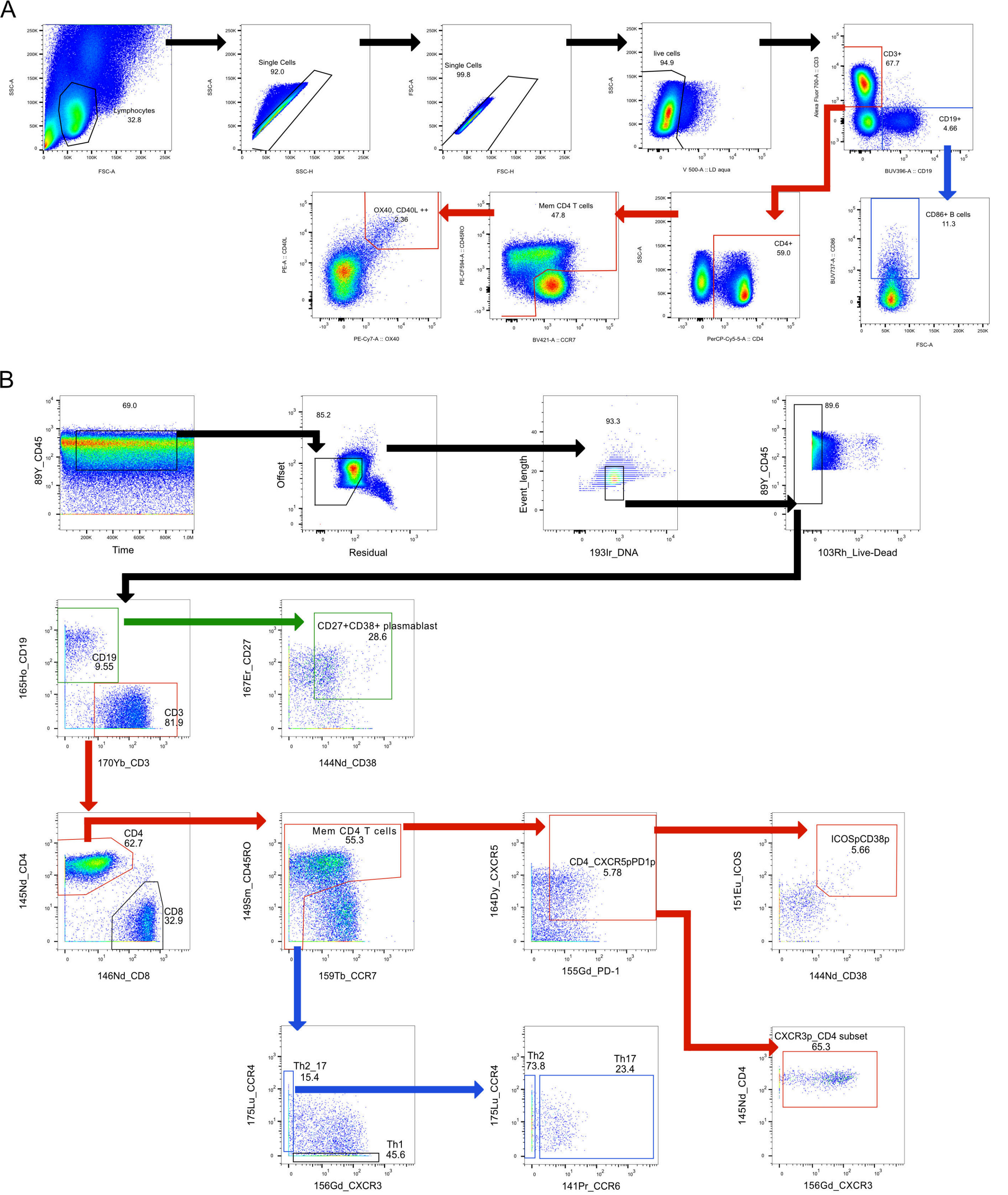

### Supplementary Figure 3

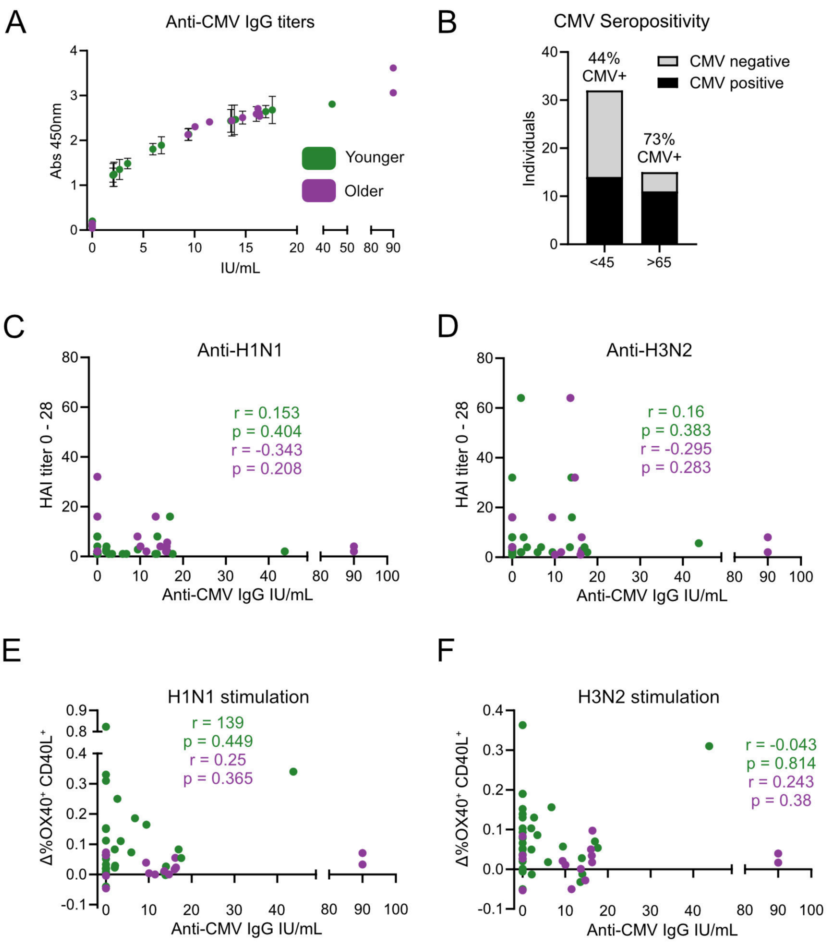

### Supplementary Figure 4

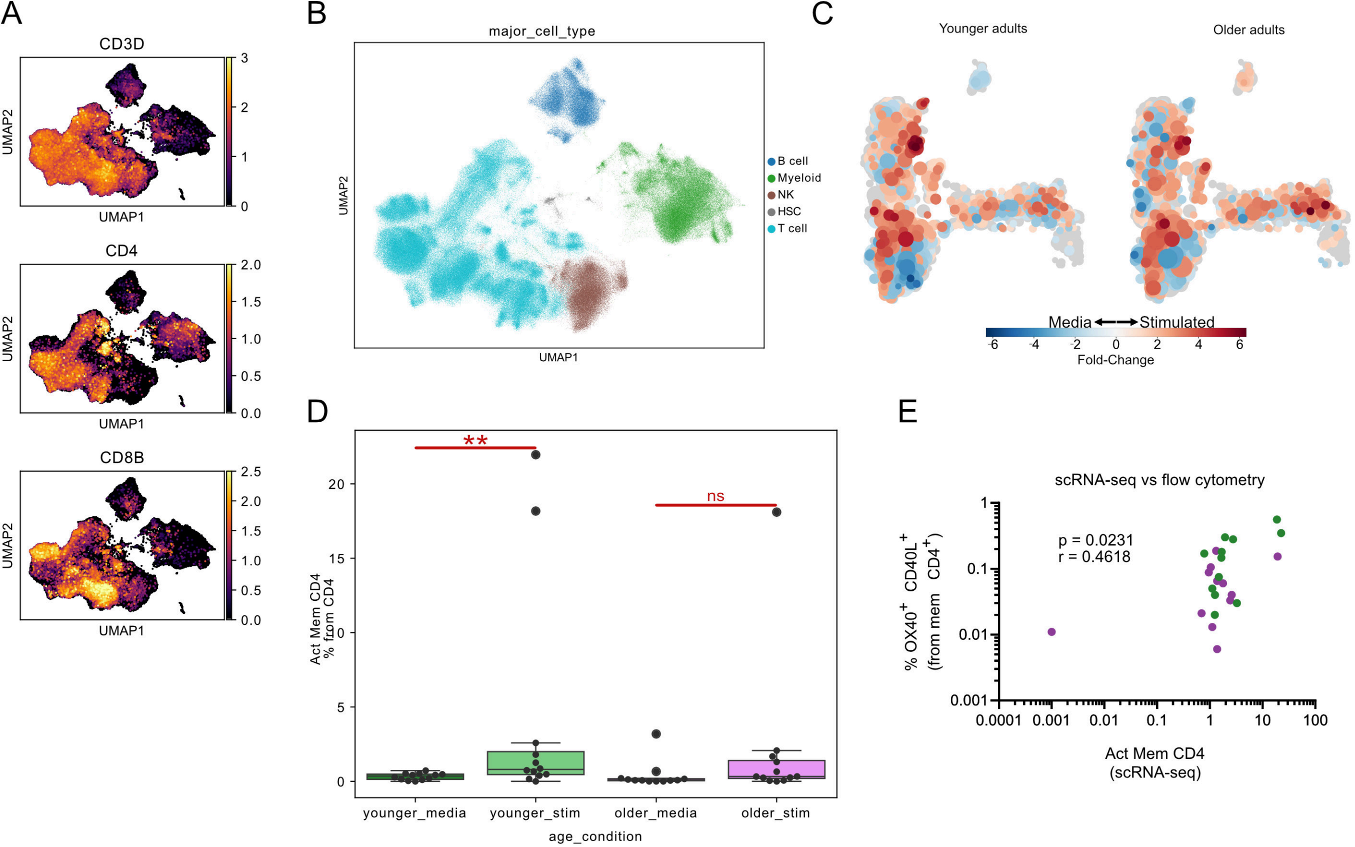

### Supplementary Figure 5

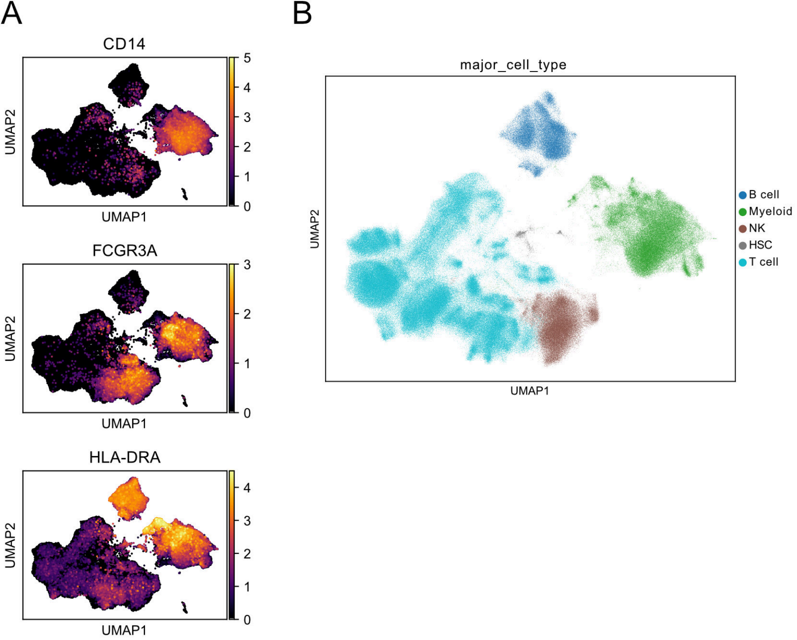
